# Modeling Chromosome Maintenance as a Property of Cell Cycle in *Saccharomyces cerevisiae*

**DOI:** 10.1101/044339

**Authors:** Jesse P. Frumkin, Biranchi N. Patra, Anthony Sevold, Kumkum Ganguly, Chaya Patel, Stephanie Yoon, Molly B. Schmid, Animesh Ray

## Abstract

Defects in DNA repair, synthesis, and chromosome transmission can often cause chromosome instability, which are understood with respect to molecular-genetic mechanisms. However, transition from descriptive models to quantitative ones is generally difficult. Here we use a computationally intensive numerical technique based on linear programming to analyze the processes of chromosome maintenance during the cell cycle in yeast, *Saccharomyces cerevisiae*. We first experimentally identify 19 genes that when ectopically expressed cause chromosome instability. We then build an 18 x 19 matrix by assaying the genetic interactions of pairs of genes that each normally functions to maintain chromosomes, including the 19 genes discovered here. We then use a ”seriation” algorithm based on linear optimization to find an optimal arrangement of rows and columns to confirm an optimum temporal arrangement of gene influence during cell cycle phases. We experimentally demonstrate that the method yields new biological insights, which we test and validate.

## Introduction

With the advent of genome-level techniques for rapid identification of gene function, it is becoming important to develop rapid methods for generating hypotheses for their mechanisms of action. One way to investigate the mechanisms by which these genes may participate jointly in a common biological process, such as chromosome maintenance or in another related biological function, is to combine loss-of-function and gain-of-function mutations in the same strain and explore their synthetic phenotype if any. For example, certain loss-of-function mutations that affect chromosome maintenance interact with other genes that affect chromosome maintenance when overexpressed.^1–4^ The phenotypes of pairs of loss-of-function and gain-of-function combinations often can reveal synergistic or suppressive effects, allowing the genes to be placed relation to one another in the context of a molecular model defining the process if at least one of the interacting partner’s molecular function is known. These genetic interactions represent n x m matrices of vectors, which contain numerical data that represent positive or negative genetic interactions between the respective genes. Such matrices embody the information contained therein, and have the potential to provide insights when they are appropriately combined with additional data.^5^ Here we investigate the method of seriation^6^ to provide insights on the cell biological mechanisms of a set of genes that affect chromosome maintenance.

The maintenance of proper chromosome copy number and stability during cell division is a highly controlled phenomenon requiring programmed expression of at least 723 genes in the yeast *Saccharomyces cerevisiae*.^7^ Some of the genes known to affect chromosome stability and maintenance function in DNA repair, replication, recombination, chromosome segregation and cell cycle control. The molecular mechanisms of chromosome stability/copy number control for the remaining genes are poorly understood, but are important for understanding mechanisms of human disorders such as birth defects, and cancers.

The involvement of a large fraction of these genes in chromosome stability and copy number control was inferred from the effects of loss of function mutation in the given gene on chromosome stability.^8–12^ Others were implicated from their adverse effects on chromosome stability when these genes were over-expressed.^13^ It is likely that more such genes remain to be discovered.

We began by discovering a set of genes whose over-expresion results in increased chromosome instability. We chose 18 of these genes and combined them with a set of deletion mutations that were known to also cause chromosome instability^10^ and assayed their combined effects on cell growth properties, anticipating that synergistic effects on chromosome stability would affect cell viability–a phenomenon termed Synthetic Dosage Lethality.^14^ The resulting matrix of genetic interactions was analyzed by integer linear programming (seriation), a relatively new technique inspired by the famous ’’Traveling Salesman Problem,” to deduce the optimal cycles among the rows and columns given the unsorted matrix of genetic interactions (minimizing the sum of 18-dimensional Euclidian distances exhibited by the 18 genes), and found one optimal arrangement among 355,687,428,095,999 alternative possible arrangements. We found that the order of the respective functions of the optimally sorted genes predicted the cyclical functions of some of these genes during cell cycle. We tested this inferred property through assessing the gene’s effects on the relative phases of the cell cycle and confirmed that the optimal order reflected their anticipated effects.

## Materials and Methods

### Strains

Chromosome stability assays and Fluorescence Activated Cell Sorting (FACS) assays were performed on cultures of the diploid strain *YPH275* of *S. cerevisiae* (*MATa/MATα ura3-52 lys2-80 ade2-101 trp1-*Δ*1 his3-*Δ*200 leu2-*Δ*1 CF [TRP1 SUP11 CEN4]*), provided by Prof. Phil Hieter.^15^ Synthetic dosage lethality and suppression were assayed by transforming individual haploid deletion mutant strains from the gene knockout strain library^16–18^ and the parental strain *BY4741* (*MATa his3*Δ*1 leu2*Δ*0 met15A0 ura3*Δ*0*) as control (available from Open Biosystems) with selected plasmids.

### Plasmids

The MORF (Movable Open Reading Frame) library consists of two-micron plasmids each having one tagged gene of interest under the control of a GAL promoter.^19^ Strains of E. coli containing the MORF plasmid library were obtained from Dr. Elizabeth Grayhack and Dr. Eric Phizicky (University of Rochester Medical School, Rochester, NY). Individual MORF clones were verified by sequencing, and were introduced into the appropriate yeast strains by transformation, and confirmed upon recovery by resequencing.

### Chemicals and Media

Chromosome instability reported by discrete red sectors on pink colonies was assayed on synthetic yeast growth media lacking uracil, which contained 5 to 6 mg per L of adenine (minimal adenine) to optimize color formation.^20^ Plates containing 2% galactose plus 1% raffinose were used to induce gene expression regulated by the GAL promoter on MORF plasmids while plates containing 2% glucose were used to repress gene expression regulated by the GAL promoter. Enriched yeast growth medium was YP with added glucose (YPD).^20^

### Transformation of MORF Plasmids into Yeast

To induce overexpression of genes, we transformed liquid cultures of cells obtained from uniformly pink colonies of *YPH275* with plasmids selected from the MORF library. *YPH275* cultures were incubated with aeration at 30°C for 2 days in 100mL of YP media including either 2% glucose (YPD) or 1% raffinose (YPRaff).^20^ Approximately 2.5 x 10^11^ cells were centrifuged into a pellet that was resuspended in 10mL of transformation buffer (0.20 M LiAc, 0.10 M dithiothreitol, 40% mass per volume of polyethylene glycol, 2.6% mass per volume of dimethyl sulfoxide, and 500 *μ*L of single-stranded DNA). Aliquots of 300 *μ*L were mixed with 5 *μ*L of plasmid DNA that was extracted using the Miniprep Kit from Qiagen. The mixture was heated for 30 minutes at 45°C and then mixed with 540 *μ*L of YPD media. 100 *μ*L aliquots were plated onto glucose-containing media lacking uracil and grown for two or three days at 30° C; the transformants were frozen in 8% dimethyl sulfoxide.

### Chromosome Stability Assay

*YPH275* transformants were single colony purified on YPD plates, followed by plating on synthetic dextrose media lacking uracil to select for the plasmids. The plasmids were resuspended in water, diluted, and plated for single colonies on media containing (1) glucose (synthetic dextrose containing minimal adenine and lacking uracil) and (2) galactose (galactose plus raffinose media containing minimal adenine and lacking uracil). After one week of growth at 30°C, plates were shifted (in multiple replicates for each strain) to 4°C and incubated for two weeks to allow red sectors to darken. In some sets of experiments, plates remained at that temperature for an additional two weeks to allow any non-specific pink color to fade without compromising the darkening of the red sectors.

The number of sectors per colony was assumed to follow a Poisson(*μ*) distribution where 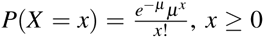. Because the sum of n Poisson(*μ*) distributions is a Poisson(n*μ*) distribution, the total number of sectors on a plate having n colonies is Poisson (n*μ*), or 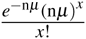. A Poisson regression to model the mutation rate per colony *μ* was performed using an offset to adjust for the number of colonies on each plate; specifically, we optimized the *β* parameter vector in log*s*_*i*_ = log *n*_*i*_ + *β*_1_*x*_1_ + *β*_2_*x*_2_ + …*β*_*n*_*x*_*n*_, where *s*_*i*_ is the number of sectors on plate i, *n*_*i*_ is the total number of colonies on plate i, and *x*_*j*_ equals one or zero to represent the *j*^*th*^ gene^*^sugar interaction.^21^ (Note that the constant term *β*_0_ was omitted from the model so that each *e*^*β*^*j* could be interpreted as the mutation rate for the *j*^*th*^ gene^*^sugar.) The significance of the differences between the estimated parameters *β*_*j*_ were computed using a Z-Test using the Z values for each parameter estimate *β*_*j*_. Significant differences between *β*_*j*_ for a gene on galactose versus the empty vector on galactose were determined by using a Bonferonni adjusted p value threshold of 0.01. The Poisson Regression was generated using SAS Software (University Edition). This technique builds on that of a dissertation,^22^ which instead reported a test of dependence of sector frequency on sugar type without using the Poisson distribution to quantify the rates of mutation per colony.

### Screening MORF Library Plasmids for Identifying Candidate Chromosome Instability Genes

Plasmid DNA comprising the entire MORF library was partitioned into sets including about 384 individual MORF plasmid DNA preparations. These DNA preparations were pooled into mixtures of 384 plasmids that were used to transform the strain YPH275 (with the expectation that only one MORF plasmid would generally be introduced into any particular transformant cell). 10*μ*1 of transformants were spotted in a dilution series on solid synthetic media plates (containing galactose, raffinose, minimal adenine, and no uracil) using a Beckman Coulter Biomek FXP liquid handler robot; *YPH275* transformed with the empty MORF vector plasmid *BG1766* was the negative control and *pGAL::CLB5* or *pGAL::YRB1* MORF plasmid was the positive control. Colonies were allowed to form, allowed to develop colony color, and dilution spots with separable colonies were examined for increased frequencies of red sectors relative to the negative control. Putative hit plasmids were extracted from the corresponding colonies, plasmid DNA confirmed by sequencing. Increased rates of plasmid instability were confirmed by retransformation into YPH275 followed by the Chromosome Stability Assay method described above.

To perform additional hypothesis-driven experiments using preselected plasmids, experiments were performed as above, but without the first step of pooling MORF plasmids and without the later step of confirming the DNA sequence of plasmids after observing frequent sectors.

### Synthetic Dosage Interaction

Individual MORF plasmids were introduced by transformation into specific haploid deletion mutant strains selected from the yeast knockout library. Transformants were selected on synthetic dextrose plates lacking uracil, and single colony purified. The BioRad SmartSpec 3000 optical density reader was used to establish equal titers among the transformants and to dilute each 10, 100, and 1000-fold. Four dilutions (1x, 10x, 100x and 1000x) were spotted in duplicate onto galactose-containing and glucose-containing solid synthetic media lacking uracil (the glucose-containing media was a control to check whether all strains at all dilutions grew equally). Three replicates of each experiment were plated and incubated at 30°C.

Growth of a given transformant in a deletion mutant strain was compared with the growth of the transformant containing the same plasmid in the wild-type background strain *BY4741* on the same plate by photographing and estimating the extent of growth at each dilution. The empty MORF vector plasmid was used as a control. Some genetic interactions were repeatedly validated to ensure consistency. When the growth of the deletion mutant strain carrying the MORF plasmid was indistinguishable from that of the wild-type background strain carrying the same MORF plasmid in all dilutions, the interaction received a score of 4. Scores of 1, 2 or 3 represented poorer growth of the mutant compared to the wild-type discernible at the various dilutions; thus, score 1 represented the most affected strain, whereas score 3 represented the least affected. The interaction score of 7 represented higher levels of growth of the mutant strain carrying the MORF plasmid compared to the wild type strain carrying the same MORF plasmid (the quantity seven was chosen so that the lowest score and the highest score are numerically equidistant from the score for normal growth).

### Seriation of Matrices of Genetic Interaction

An unsorted matrix *M* was built using 19 row vectors of MORF plasmids, including the empty vector plasmid, and 18 column vectors of deletion mutants. As noted above, we used a score of 1 to represent synthetic dosage lethality, 2 or 3 to represent levels of synthetic dosage sickness, 4 to represent normal growth or that data were not available, and 7 to represent suppression. Given the matrix of genetic interactions M, it is computationally intensive (and NP hard) to find the one optimal arrangement of the 19 row vectors of M that minimizes the sum over all of the 18-dimensional Euclidean distances between the rows of data vectors.^23^ Nevertheless, the optimal rearrangement of the row vectors that minimizes this sum of 18-dimensional Euclidean distances can be solved using integer linear programming^24, 25^ because the total number of row vectors in our work is sufficiently small.

This optimization task is similar to the famous ’’Traveling Salesman Problem”, except that the two-dimensional Euclidean distance between two cities in the traditional Traveling Salesman Problem is replaced here by the 18-dimensional Euclidean distance between the data vectors of scores in rows i and j of our matrix M. Specifically, if we denote our 18-dimensional Euclidean distances *d*_ij_, our optimization task is to find a vector *x*_ij_ comprised of zeros and ones (where one represents the adjacency between rows i and j), while minimizing *D*(*x*) = ∑_*i*> *j*_ *d*_ij_*x*_ij_ subject to the constraints that 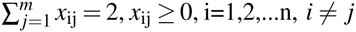 and *x*_ij_ = *x*_ij_. This optimization problem was solved using Mathematica Version 10,^26^ by using the ”FindShortestTour” function, specifying the “Method” ”IntegerLinearProgramming” and the ”Distance Function” of ”EuclideanDistance”.

To sort the columns in addition to the rows, the resulting matrix was transposed and ”FindShortestTour” was used to re-sort the transposed matrix using the same method and distance function.

### Fluorescence Activated Cell Sorting Analysis

To measure the effects of MORF plasmids on the percentage of cells in G1, we prepared cultures of *YPH275* for scanning by Imaging Flow Cytometry.^27^ Transformants were grown at 30°C in 96-well plates that contained 200 *μ*L of synthetic dextrose media lacking uracil. After two days of growth, 10 *μ*L of cells were inoculated into fresh 96-well plates and grown overnight in synthetic complete media with raffinose and lacking uracil. For 16 hours at 30°C, the *pGAL* promoters were induced using synthetic complete media with 2% galactose and lacking uracil, and cell density was monitored to obtain cultures at the exponential growth phase. Approximately 2 x 10^6^ cells at the mid exponential phase were harvested by centrifugation and fixed using 70% ethanol by gently mixing at 4°C. The fixed cells were collected by centrifugation, which were then washed in water. The cells were treated with 1 mg/ml RNAse A (Sigma) for 4 hours at 37°C. After centrifugation, the pellets were incubated with 1 mg/ml of Proteinase K (Invitrogen) for 1 hour at 50°C. Cells were collected by centrifugation and re-suspended with 1 *μ*M of SYTOX green (Invitrogen) in 50mM Tris (pH 7.5).

Cells were scanned by an imaging flow cytometer, Amnis ImageStreamX (Amnis Inc., Seattle, WA). 25,000 cell images were acquired and the intensity of the DNA image (green channel; 480-560nm) was recorded along with the aspect ratio of the cell image (brightfield). The Mathematica^26^ function ”FindClusters” was used to compute four clusters based on standardized Z-scores of the intensity and aspect ratio in each experiment (where each Z-Score is 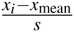, where *x*_*i*_ is a raw observation, *x*_mean_ is the mean of those raw observations, and s is the standard deviations of those raw observations. Compared to the other clusters, the cluster for G1 was assumed to have the high aspect ratio and low DNA intensity. The percentage of cells in G1 was estimated by dividing the number of cells in the G1 cluster by the total number of cells. Note that this technique builds on that of a dissertation^22^ which instead assumed predetermined thresholds of DNA intensity and aspect ratio to demarcate cells in G1.

## Results

### Identifying Genes Causing Chromosome Instability Upon Overexpression

The yeast strain *YPH275* is a diploid that is homozygous for the *ade2-101* ochre mutation, and also contains the dominant ochre-suppressor gene *SUP11* on a supernumerary chromosome fragment.^15^ The loss of the supernumerary chromosome or the *SUP11* gene is signaled by the accumulation of a red pigment in the colony (or colony sector) due to defective adenine biosynthesis in the absence of *SUP11*. In principle, the loss of *SUP11* can also result from ectopic gene conversion to *sup11* present at the allelic locus, but this frequency is generally low compared to the frequency of loss of *SUP11* by mitotic nondysjunction or recombination. The increased frequency of red colonies/sectors on galactose relative to the background levels on glucose is a measure of chromosome instability induced by the MORF plasmids.^28^

With the expectation that many genes that affect the stability and copy number of chromosomes when overexpressed remain to be identified, we took two independent approaches, which are described below.

1) In a model independent approach, we screened the entire Movable Open Reading Frame library (MORF) using a technique^13^ having a high false negative rate; using this technique Ouspenski and colleagues discovered 30 genes that induce chromosome instability upon over-expression but had estimated that 58 genes had remained to be discovered. Using this screen technique followed by validation, we confirmed eight MORF plasmids as capable of inducing chromosome instability when over-expressed on galactose, including *XRS2*, *HFI1*, *ELG1*, *CLN1*, *GRX2*, *NOP6*, *SPC19*, and *GPG1* (we also confirmed the positive control *YRB1*). Using the same assumptions as Ouspenski [13, see p. 3002], we estimate to have missed at least 40 additional genes using this technique. These genes belong to pathways known to participate in chromosome maintenance, DNA replication, recombination, repair, and cell cycle progression.
(2) Our second approach was an intuition-dependent approach. We had noted the possibility that abnormal expression of meiosis related genes in mitotic cells might cause mitotic chromosome instability. To explore this further, we chose 25 meiosis-related genes (*DMC1*, *DON1*, *GIP1*, *HFM1*, *HOP1*, *HOP2*, *MCK1*, *MEI5*, *MEK1*, *MER1*, *NDJ1*, *NDT80*, *REC8*, *SAE3*, *SET3*, *SPO1*, *SPO11*, *SPO13*, *SPO19*, *SPO20*, *SPO22*, *SPO73*, *SPO75*, *SPO77*, and *ZIP2*) and tested these for galactose induced chromosome instability. Only two of these genes, *NDT80* and *REC8*, induced chromosome instability in our assays (*i.e*, a hit-rate of 8%. Predominantly red sectors, characteristic of the clonal loss of the supernumerary chromosome, against a light pink colony background (characteristic of the ochre-suppressed *ade2-101* phenotype) that are unaccompanied by white sectors (characteristic of cells with additional copies of the supernumerary chromosome) occurred in strains containing *pGAL::NDT80* and *pGAL::REC8*. This observation indicated that mostly chromosome loss were induced by these genes without non-disjunction, or that hyperploidy and/or *SUP11* dosage might be toxic to those strains, causing selective loss of aneuploid cells.

In addition to testing meiosis-specific genes, we also tested various genes that function in mitotic cell cycle control and genes that had been previously assayed during a screen for dosage suppression.^29^ We screened 295 of these plasmids and retested 40 of these to confirm 17 additional genes that induce chromosome instability (about 6% of the 295 screened). The confirmed genes include *BMT5*, *CDC4*, *CDC5*, *CDC20*, *CDH1*, *CWC24*, *HMLα2*, *IME2*, *MCM3*, *MOT2*, *NRM1*, *SLD3*, *SLM4*, *SPO7*, *YDR387C*, *YGL182C*, and *YLR053C*. (Note that *HMLα2* is ordinarily silenced, but is regulated by an inducible *pGAL* promoter in the MORF library.) In addition, we reconfirmed the positive control *CLB5*.

We quantified the chromosome loss rate per colony (*μ*_*coiony*_) induced by the MORF plasmids relative to that by the empty vector *BG1766* by Poisson regression as described in Methods. Table 1 shows the results of these computations. The sets 2, 5, 7, 8, 9, 10, 11 and 12 include genes identified by the chromosome instability screen described above that was similar to that of Ouspenski et al. Set 4 includes genes identified by screening genes that normally function during meiosis. The sets 1, 3, and 6 include genes that were chosen because of their role in cell cycle control or because they were assayed during a screen for dosage suppression. In view of the results in set 6, we propose naming the uncharacterized ORF *YDR387C* to *CIN10*.

**Table 1.**
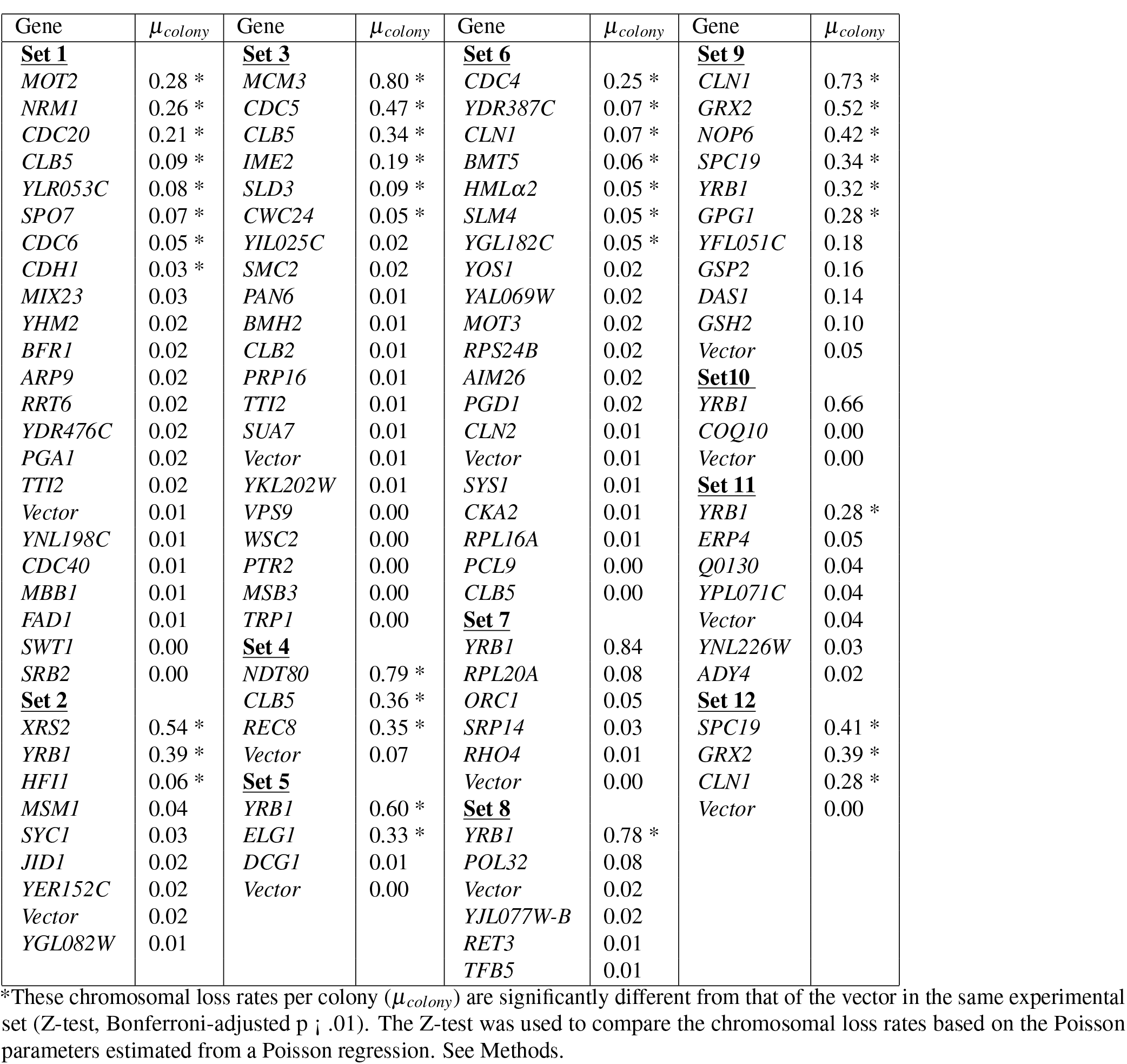
The Rate of Chromosomal Loss per Colony (*μ*_*colony*_) by Conditional Overexpression Using Diverse Sets of Genes

Note that some sector-inducing genes are not reported because red sectors were difficult to distinguish visually from pink sectors. The excluded results are those of *pGAL::YML007-C, pGAL::YAR023C, pGAL::MES1*, and *pGAL::MHT1*. In addition, one batch, Set 7, the positive control had a small number of colonies plated and hence an insignificant mutation rate per colony.

### Seriation of a Matrix of Genetic Interactions to Form a Genetic Model for Chromosome Instability Induction

We hypothesized that some of the genes might be affecting chromosome stability due to their adverse effects on the cell cycle. If true, it is predicted that these genes should affect relative lengths of the cell cycle in a predictable manner. One way to numerically identify whether a set of vectors have systematic effects on a cyclically timed series of events is to conduct seriation analysis on the matrix of vectors. Based on the *μ*_*colony*_ estimated as described in the above section, 19 plasmids were chosen for matrix seriation analysis. The corresponding genes are *CDC4*, *CLN1*, *ELG1*, *YGL182C*, *NRM1*, *NDT80*, *IME2*, *REC8*, *CDC5*, *YRB1*, *CLB5*, *MOT3*, *CDC20*, *XRS2*, *YDR387C*, *MOT2*, *HMLα2*, *CSE4* and *HHT2* (of which *CSE4* and *HHT2* were included despite undetermined mutation rates). These 19 MORF plasmids were introduced into 18 deletion mutant strains, each of which gene deletions were previously reported to cause defective chromosome maintenance [10, Supplementary Table S3]. The resulting 331 combinations (11 strains could not be constructed for technical reasons) were assayed for synthetic growth defect or suppression of growth defects as described in Methods. The resulting matrix of suppression, synthetic dosage lethality, and synthetic dosage sickness is illustrated in Figure 1. Not shown are the rows for *pGAL::HHT2, pGAL::CDC4, pGAL::CLN1*, and *pGAL::HMLα2*, which are identical to the row for the empty vector. Note that this matrix has no dimension of time: the possibility that the MORF plasmids might be exhibiting the genetic interactions observed here might have to do with any temporally cyclical phenomenon (such as the cell cycle) is implicit. The matrix was seriated by Integer Linear Programming (see Methods) for optimizing the total of the Euclidean-distance metrics between pairs of row vectors. The optimum sequence of the rows in the matrix had a total space of 355,687,428,095,999 permutations. The observed optimal order of the MORF genes following seriation is given in Figure 1. When we listed the genes Gene Ontology annotations^30^ of the deletion mutant genes, we recognized that the observed series form a kinetically coherent model approximating the temporal order of cell biological events that occur during cell cycle:

1. DNA damage sensing (e.g., *TOF1, RAD9, MEC3*);
2. Kinetochore-centromere cohesion (e.g., *IML3, CHL1*);
3. Helicase activity (e.g., *CHL1*);
4. Spindle checkpoint delay (e.g., *MAD2*);
5. DNA stress response (e.g., *SPT21*; *DDC1*, *MMS22*, *RAD54*);
6. Attachment of kinetochore to microtubules (e.g., *MCM21*);
7. Initiation of anaphase (e.g., *MAD1*);
8. The progression through the spindle assembly checkpoint (e.g., *CHL4*);
9. Chromosome segregation (e.g., *MCM22*);
10. Inhibition of cohesion between sister chromatids (e.g., *RAD61*);
11. Re-linking sister chromatid cohesion to DNA synthesis (e.g., *CTF4*). This function is followed cyclically by DNA damage sensing again.

**Figure 1.**
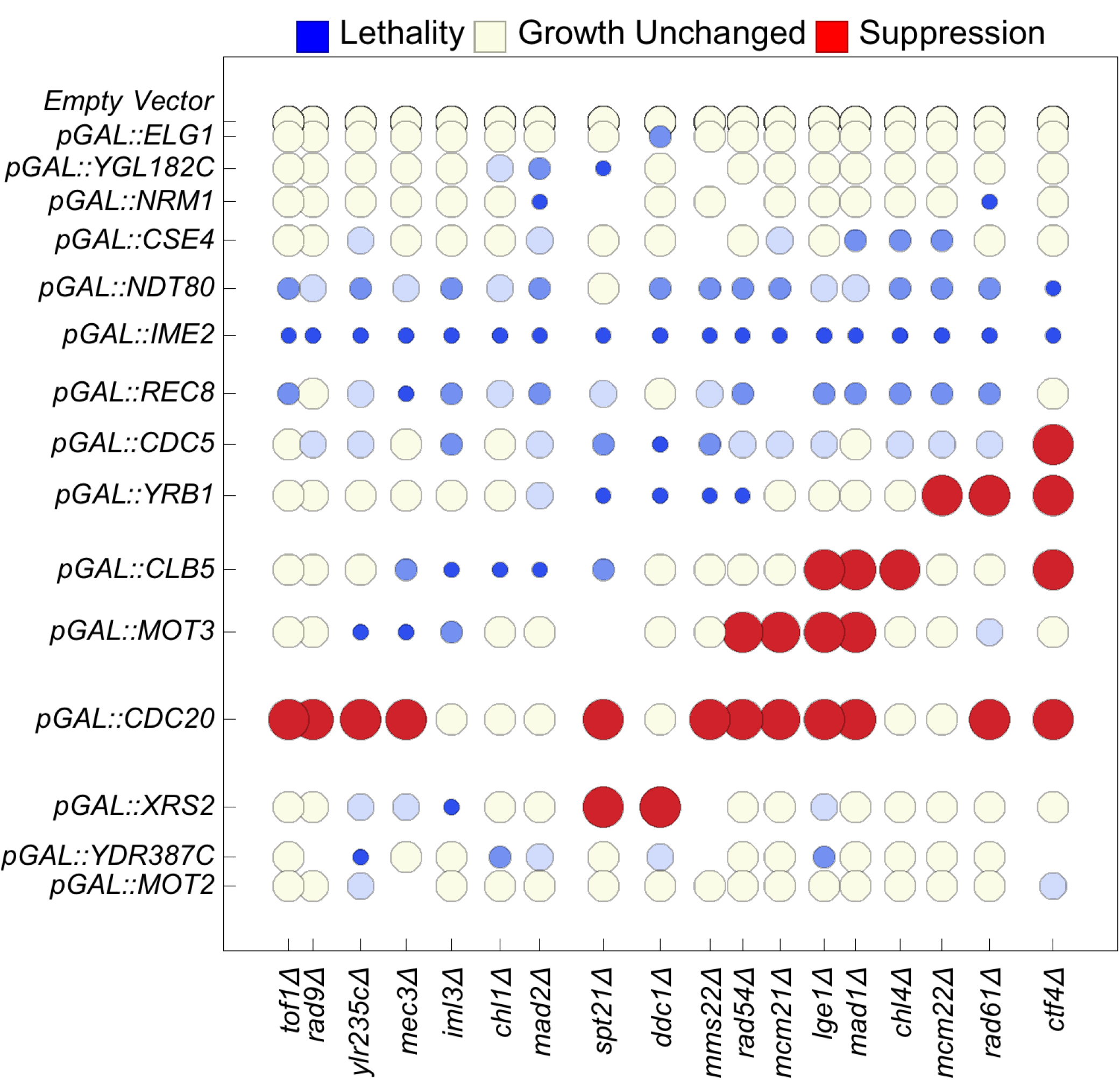
Seriation of genetic interactions results in a cyclical matrix. We report synthetic dosage lethality (represented as small blue circles), two levels of synthetic dosage sickness (two progressively lighter shades of blue), unchanged growth (yellow), and suppression (large red circles). To seriate the matrix, the rows (columns) of the matrix of genetic interactions are rearranged using integer linear programming to minimize the sum of the n-dimensional Euclidean distances over all pairs of adjacent n-dimensional row vectors (column vectors). The spatial distances of separation between adjacent rows and adjacent columns are proportional to the n-dimensional Euclidean distance between two separated rows or columns, respectively. The resulting order of the genes forms a model of the order of the respective functions of the genes during the cell cycle. The first row (column) is adjacent to the last row (column). Not shown are the rows for *pGAL::HHT2*, *pGAL::CDC4*, *pGAL::CLN1*, and *pGAL::HMLα2*, which are identical to the row for the empty vector.

We predict that one gene of unknown function, *LGE1*, has a similar function to the cyclically adjacent genes *MCM21* (functioning in kinetochore attachment) and *MAD1* (functioning in the spindle checkpoint). We predict that the dubious open reading frame *YLR235C* is similar to the cyclically adjacent DNA damage sensing genes *RAD9* and *MEC3*, perhaps because the deletion of *YLR235C* overlaps with the deletion of *TOP3* (topoisomerase III).

To arrange the overexpressed genes and the respective annotations^30^ of those genes, we again minimized the total of the n-dimensional Euclidean distances rather than maximizing that total. The model arrangement is:

1. Regulation of the G1 to S transition (e.g., *CDC4, CLN1*);
2. DNA replication and repair (*e.g., ELG1*);
3. Exit from G1 (*e.g., NRM1*)
4. Kinetochore function (e.g., *CSE4*)
5. Positive regulation of meiosis (e.g., *NDT80, IME2*)
6. Sister chromatid cohesion during meiosis (e.g., *REC8*)
7. Various functions during meiosis or mitosis (e.g., *CDC5*)
8. G1 to S transition (e.g., *YRB1*)
9. DNA synthesis during S phase (e.g., *CLB5*)
10. Regulation of stress response (e.g., *MOT3*)
11. Finish stage of Mitosis (e.g., *CDC20*)
12. Double Strand Break Repair (e.g., *XRS2*)
13. Regulation of DNA replication (e.g., *MOT2). MOT2* is followed cyclically by G1 functioning genes again.

This second sequence is similar to the first sequence above. Each predicts that DNA repair is followed by kinetochore function, followed by stress response, followed by completion of mitosis, followed by DNA synthesis. This second model includes an excursion into meiosis after exiting G1 because meiosis-specific genes were used to build the matrix.

### Evaluating the Model Using Fluorescence-Activated Cell Sorting

Does the order of the overexpressed genes in the model correctly predict the order of the respective functions of the genes during the cell cycle? To validate the model, we tested whether genes that function to transition the cell cycle out of G1 reduces the percentage of cells in G1,^31^ while genes that function to transition the cell cycle out of M Phase do the opposite. The percentage of cells in G1 are inferred by clustering two-dimensional data consisting of (1) normalized DNA fluorescence and (2) normalized aspect ratio of the brightfield image. Of importance to note here is that chromosome instability induced by plasmid overexpression was conducted in the diploid strain *YPH275*, whereas the temporal sequence of events were inferred from the synthetic genetic interaction matrix based on cell viability in haploid cells. Therefore the model validation experiments by FACS were conducted in the diploid *YPH275* background.

Figure 2 shows the percentage of diploid cells in the G1 phase of the cell cycle for *YPH275* strains carrying the respective MORF plasmids, when grown in the presence of galactose. The order of the genes in this figure is sorted to match the order of the cycle in Figure 1. Recall that the genetic interaction data for *CDC4*, *CLN1*, and *HMLα2* that are reported Figure 1 are identical to that of the empty vector. Data are not shown for *AIM26*, a gene of unknown function that was not tested for genetic interactions. With *AIM26*, we observed that the percentage of cells in G1 was 10%, 10%, and 11%. Data for *pGAL::NRM1* are excluded due to technical difficulties.

**Figure 2.**
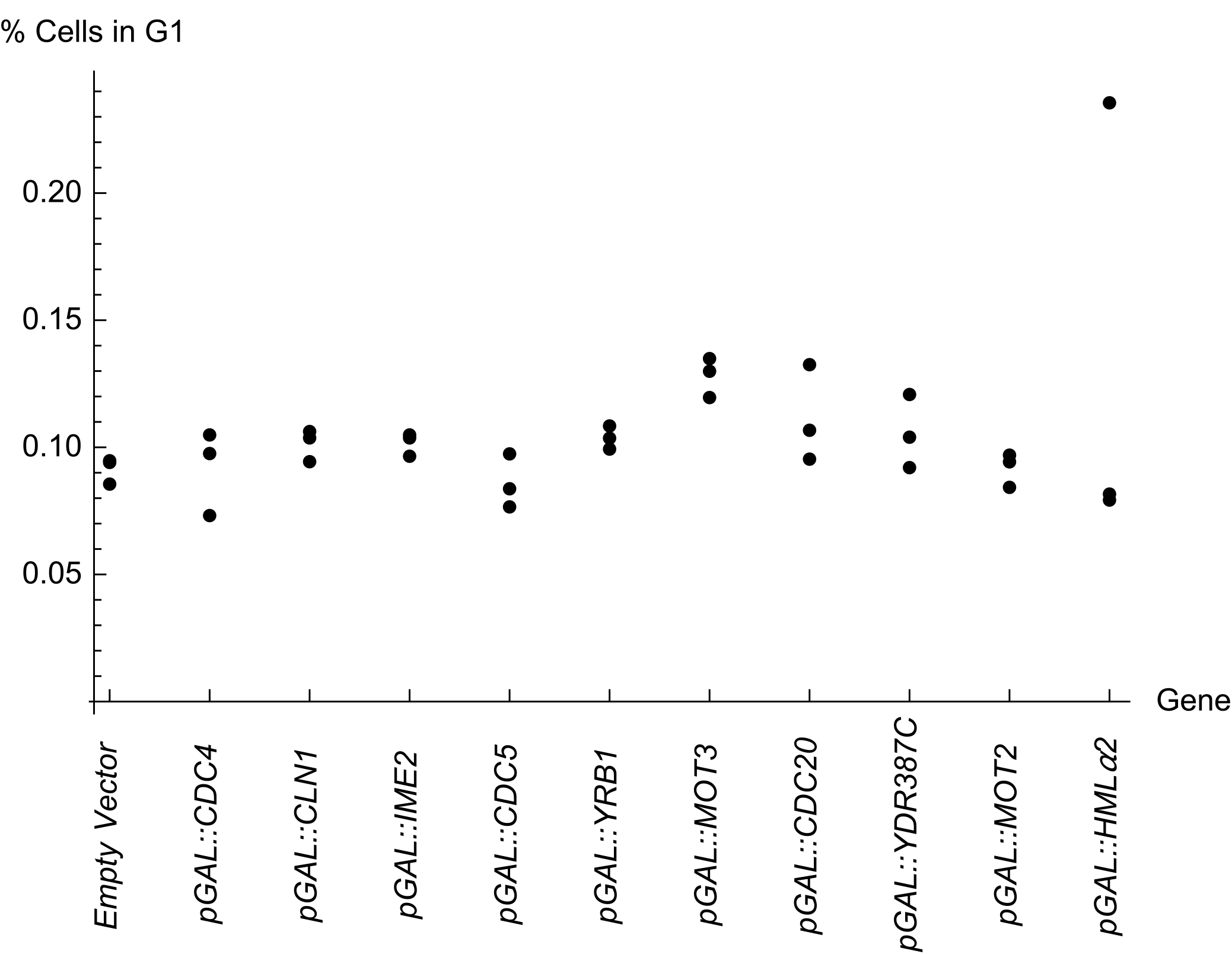
The percentage of cells in G1 measured by Fluoresence Activated Cell Sorting (FACS). The arrangement peaks with *pGAL::MOT3*. The order of genes is arranged to match the order of the cycle in Figure 1. Recall from Figure 1 that *pGAL::CDC4*, *pGAL::CLN1* and *pGAL::HMLα2* had no genetic interactions and are thus similar to the empty vector. Data for *pGAL::NRM1* are excluded because the results were irreprodicible among replicates.

Encouragingly we see further evidence of a cycle. The arrangement from *BG1766* to *pGAL::IME2* to *pGAL::CDC5* to *pGAL::YRB1* to *pGAL::MOT3* progresses toward a peak significantly (p=0.0062, Jonckheere-Terpstra Test, a non-parametric test for ordered differences among classes). Furthermore, the arrangement from *pGAL::MOT3* to *pGAL::CDC20* to *pGAL::YDR387C* to *pGAL::MOT2* to *BG1766* declines from that peak significantly (p=0.0024, Jonckheere-Terpstra Test). Hence, the arrangement supports a cyclically ordered model. Other overexpressed genes, including *pGAL::CDC4, pGAL::CLN1*, and *pGAL::HMLα2* are not implicated in genetic interactions in this work and, therefore, it is reasonable that those percentages of cell in G1 do not appear to be trended (assuming that the data for *pGAL::HMLα2* includes one outlier point).

We observed that *pGAL::CDC20*, induces a high percentage of cells in G1, while *pGAL::CLB2* induces one of the lowest percentage of cells in G1 (data for *pGAL::CLB2* that is not in Figure 2: 8%, 8.5%, and 8.4%, respectively, in three replicate experiments). These results are intuitively understandable given previous observations that *CDC20* functions in a manner opposite to that of CLB2.^32, 33^

### Seriation of a Matrix of *in silico* Genetic Interactions Involving *CDC20* and *CLB2*

The mitotic spindle checkpoint ensures that chromosomes are properly aligned and oriented in preparation for the metaphase-to-anaphase transition. Loss-of-function mutations of the spindle checkpoint genes *mps1*, *mad1*, *mad2*, *mad3*, *bub1*, or *bub3*, all of which affect this process, cause chromosome instability.^10,34–37^ In the presence of a functional M-phase spindle checkpoint, cells will usually undergo proper cell division but do occasionally bypass checkpoint arrest and undergo premature chromosome segregation. This latter effect has been called “mitotic slippage” or “adaptation”, and can occur after periods of cell cycle checkpoint arrest.^38^ The consequences of one generation of mitotic slippage may often include aneuploidy and inviability.^39,40^ The spindle checkpoint genes noted above encode proteins that regulate the sequestration and subsequent release of Cdc20p to trigger the metaphase-to-anaphase transition at the appropriate time.^41–43^ Since such mechanisms are generally thought to be important for chromosome copy number stability and genetic interactions, we used a system of coupled linear differential equations to model cell cycle events,^32^ and systematically varied certain parameters for behavior of the numerical solutions that could simulate irregularities in cell cycle oscillations and thus might predict chromosome instability and dosage suppression.

In the model, protein concentration, phosphorylation, protein complex formation, etc., of the minimal components of cell cycle regulation are modeled using kinetic rate laws combined with algebraic constraints and conditional logic as described previously by Chen and her colleagues.^32^ Alteration of parameters in this mathematical model can simulate gene deletion, gene dosage effects, binding affinity changes, and several other perturbations.^32^ For example, setting the rate constant for Cdc20p basal synthesis 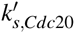 to a higher value simulates a higher level of CDC20 gene expression driven by the *GAL* promoter:

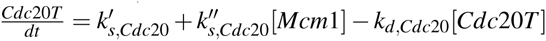

where [*Cdc20T*] is the total concentration of active and inactive Cdc20p, 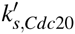 is the rate constant for basal synthesis, 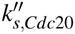 is the rate constant for Mcm1p-dependent synthesis of Cdc20p, [Mcm1] is the concentration of Mcm1p protein, and *k*_*d*,*Cdc*20_ is the rate constant for Cdc20p degradation. The reasons for using these exact variables were defined previously.^32^

Cell cycle arrest was numerically simulated in the model in a systematic manner, resulting in loss of periodicity of protein concentrations and loss of execution of a proper sequence of cell cycle milestone events [32, see the “robustness criteria”]. Upon mathematically generating cell cycle blocks through a primary perturbation, we systematically searched the parameter space in the system of equations to identify secondary perturbations that would allow resumption of the cell cycle. See Figure 3. Changes of values in parameters that would cause the simulated cell cycles to resume their periodic behavior were noted. For the purpose of this investigation, we focused on the simulated gain-of-function perturbations.

**Figure 3.**
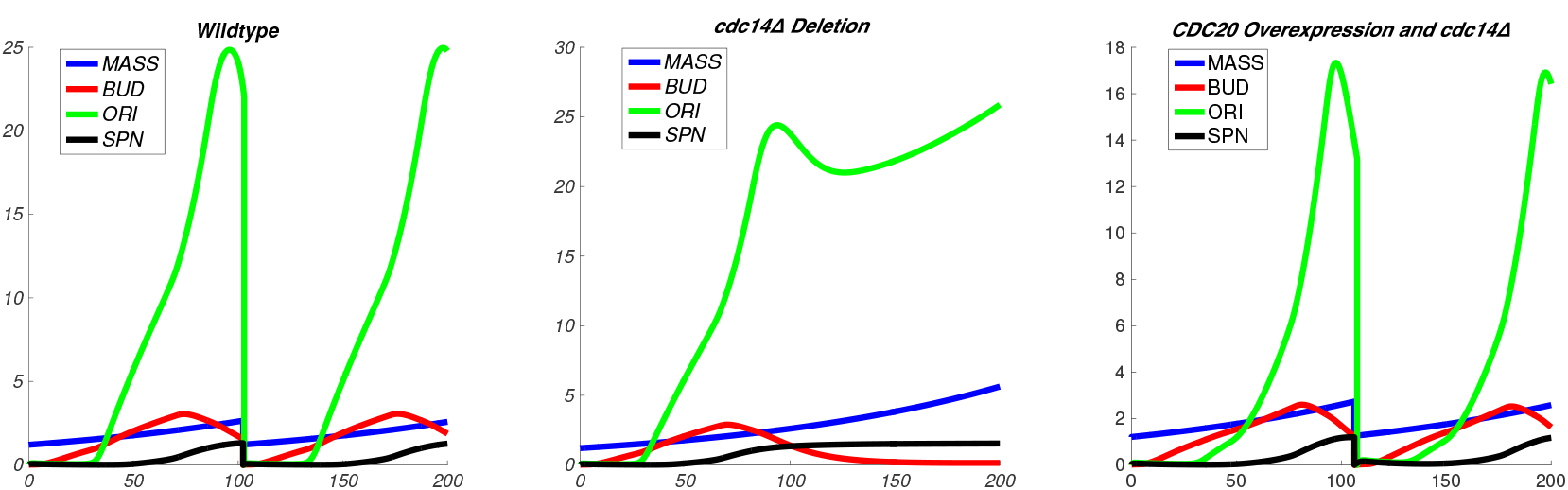
Simulations of the timecourse of *in silico* cellular mass (MASS), budding (BUD), development of origins of replication (ORI) and spindle formation (SPN) in a wildtype model^32^ (left), *cdc14*Δ deletion model (center) and a model with *CDC20* overexpression in a *cdc14*Δ deletion background. Deletion of *cdc14*Δ was simulated by zeroing the parameter representing the basal synthesis rate of Cdc14 protein, while *CDC20* overexpression was simulated by increasing the basal synthesis rate of *CDC20* from 0.006 to 1.5 dimensionless units.

In this manner, we identified five genes that bypassed the arrest induced by loss (or “in silico mutation”) of genes that act downstream from the checkpoint signaling genes. (Conceptually, the suppression in model effectively restores the signal that the checkpoint condition has been satisfied.) Specifically, the model predicts that *CDC15* should suppress a downstream *cdc14* mutation; *TEM1* over-expression should suppress a downstream *cdc15* or *cdc14* mutation; and either *CDC20* or *CDH1* overexpression should suppress a downstream *tem1, cdc15*, or *cdc14* mutation, etc.

If the seriation approach is a valid way to deduce cyclical models from genetic interactions, then we expect that seriation of the genetic interaction data should reproduce the underlying qualitative assumptions about the progression of events in the predetermined cell cycle model. For example, in the model Tem1p reacts with Cdc15p, which reacts with Cdc14p, which reacts with Cdc20p; therefore we expect that seriation should result in a similar ordering. Encouragingly, we computed that seriation of the matrix of the *in silico* genetic interactions results in the arrangement of genes that closely matches the order of the respective function of the genes in the model (see Figure 4).

**Figure 4.**
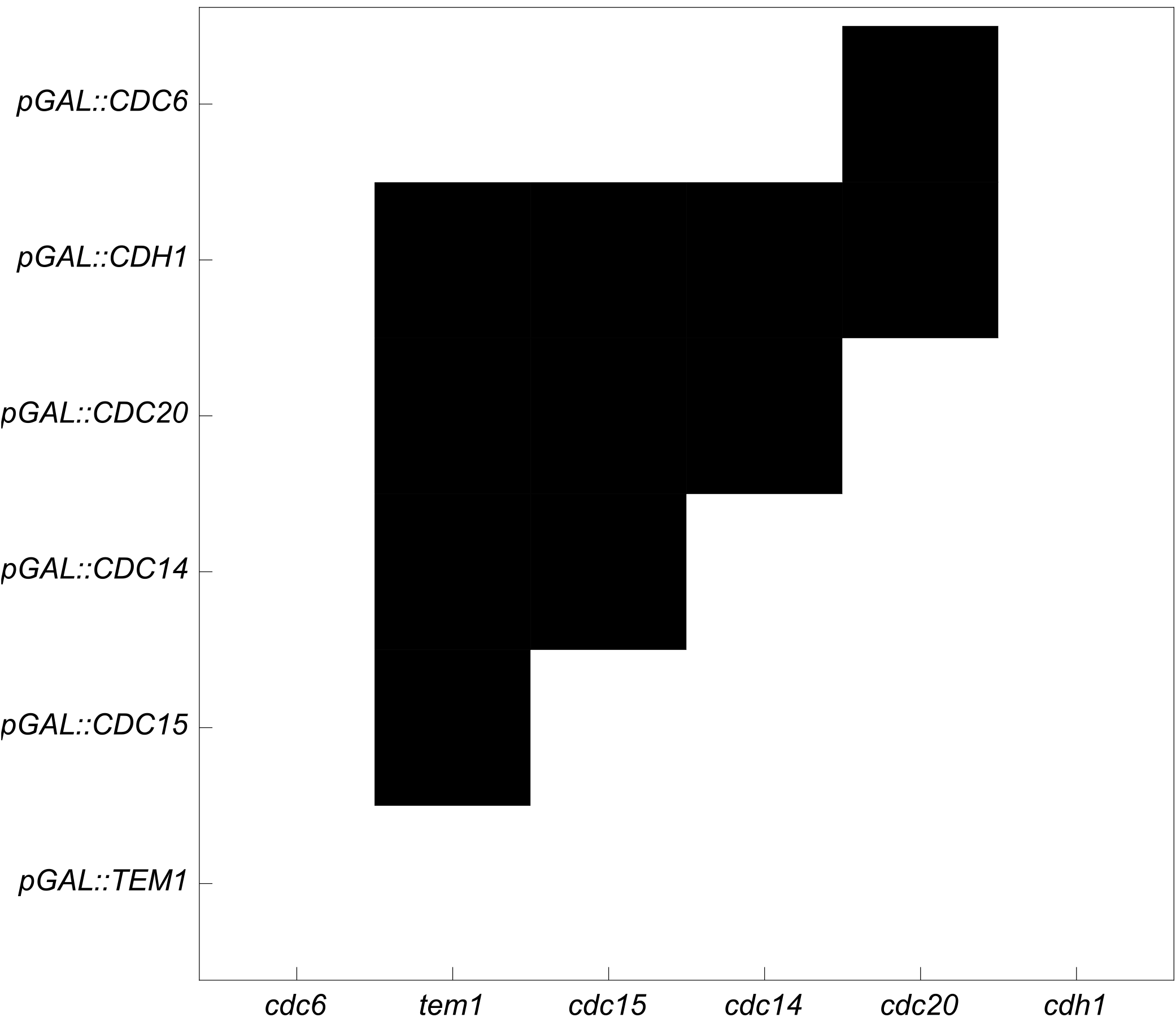
Seriation of *in silico* genetic interactions results in a cyclical matrix where the first row (column) is adjacent to the last row (column). We represent the *in silico* dosage suppression as black cells in the matrix. To seriate the matrix, the rows (columns) of the matrix of genetic interactions are rearranged using integer linear programming to minimize the sum of the n-dimensional Euclidean distances over all pairs of adjacent n-dimensional row vectors (column vectors). The predicted order of the genes reproduces the ordering of events in the underlying model that was used to simulate the dosage suppression (e.g., Tem1p reacts with Cdc15p, which reacts with Cdc14p, which reacts with Cdc20p).

## Discussion

The 34 phenotypes of synthetic dosage lethality, 38 of synthetic dosage sickness, and 26 of suppression are novel according to BioGrid (www.thebiogrid.org). Our overexpression-induced sectoring phenotypes are novel in *Saccharomyces cerevisiae* as well, other than the positive controls *YRB1*,^13^ and *CLB5*.^28^ Nevertheless, mechanisms that might suggest chromosome instability have been explored by others, including works that study *CDC5* overexpression^44,45^ and *CDC20* overexpression.^43,46–49^

Suppression was most pronounced for combinations that included *CDC20:* 12 out of 18 combinations suppressed the deleterious effects of *CDC20* overexpression. This is intuitive because *CDC20* causes increased white sectors indicative of a chromosome gain or gene copy number gain (reported in this work). On the other hand, the various deletion mutations each is known to cause increased frequency of chromosome loss.^10^ Thus, the suppression phenotype can be simply explained by mutually compensatory effects of the respective genetic perturbations.

In this work, *pGAL::MOT3* also caused frequent chromosome gain events as signaled by the occurrence of predominantly white sectors even in the absence of adjacent red sectors. Hence, colonies overexpressing *pGAL::MOT3* appeared similar to *pGAL::CDC20*. Indeed *CDC20* and *MOT3* each cause a high shift to the percentage of cells in G1 (this work).

*MOT3* encodes a prion that affects Ty transposable elements,^50^ yet *pGAL::MOT3* was not previously implicated in chromosome instability until this work. Nevertheless, it has been reported that loss of *spt4* function affects Ty transposable elements^51^ and also induces chromosomal instability.^52^ Furthermore, *spt4* loss of function can increase the growth of yeast cells.^53^ Hence, transposable elements may be one of the few genetic elements affecting both chromosome instability and increased growth, both of which are both hallmarks of cancer.

The white-sectored colonies, observed using *pGAL::CDC20* and *pGAL::MOT3*, could result from *SUP11* expression. The *SUP11* ochre suppressor gene, can increase the rate of spontaneous nondisjunction during meiosis.^54^ Unlike the suppression observed using *pGAL::CDC20*, there are the extensive synthetic dosage lethal interactions between the deletion mutations and three meiosis-specific genes, *IME2*, *NDT80*, and *REC8*.

*IME2* encodes a meiosis-specific serine-threonine protein kinase that activates the early stage of meiosis. The activation triggers the transcription factor Ndt80p; the latter is a master controller of middle-sporulation genes and is a member of the transcription factor family that includes p53.^55^ Overexpression of *IME2* in haploid cells reduce cell division rate, possibly due to a G2/M phase delay.^31^ The observation that all 18 deletion mutant strains in combination with *IME2* overexpression exhibit synthetic dosage lethality may indicate a non-specific toxicity of Ime2p. Alternatively, the effect of *IME2* overexpression could be due to a specific effect on chromosome dynamics that is enhanced in combination with each of the 18 deletion mutations that are also known to affect chromosome dynamics. Similarly, with *NDT80*, synthetic dosage lethality interactions were obtained using 17 out of 18 deletion mutants tested (the strongest was with *ctf4*Δ).

No direct genetic or physical interactions were reported previously between *REC8*, a meiosis-specific protein required for maintaining sister-chromatid cohesion and preventing sister-chromatid recombination, and *TOF1*, a gene necessary for sister-chromatid cohesion following DNA damage. The synthetic dosage lethal interaction reported here between *REC8* and *tof1*Δ establishes this connection, consistent with the idea that both genes are part of a single mechanism that controls sisterchromatid cohesion. Similarly, *MEC3* encodes a sliding clamp protein for replication fork movement, with no reported direct physical or genetic interaction with *REC8* or *TOF1*. The synthetic dosage lethal interaction reported here between *REC8* and *mec3*Δ illustrates a further mechanistic link between replication fork movement and sister-chromatid cohesion/recombination.

Paradoxically, overexpression of *NDT80* lengthens the replicative lifespan of yeast,^56^ despite that *pGAL::NDT80* is toxic upon overexpression using glycerol-containing media.^19^ *REC8* overexpression also affects the replicative lifespan^57^ and yet induces sectors (this work). Note that the red sectors observed using *NDT80* and *REC8* in the work are not caused by nondisjunction because no white sectors were observed.

Suppression and synthetic dosage lethality among several genes helped us establish novel functional connections between pairs of genes. For example, *CDC5*, which encodes a polo-kinase important for several functions in mitosis including DNA damage-repair, exhibits strong synthetic dosage lethality with the *ddc1*Δ, which encodes a DNA damage checkpoint protein. The synthetic dosage lethality is understandable if one assumes that excess Cdc5p activity exhibits dominant negative effects on DNA-damage repair, which would be expected to interact synergistically with *ddc1*Δ mutation. This theory is supported by the observation that *CDC5* overexpression alone inhibits growth, which is suppressed by *ctf4*Δ, which is required for sister-chromatid cohesion at repair-replication forks.

*XRS2*, which encodes a protein essential for nonhomologous end joining at DNA double-strand breaks, exhibits synthetic dosage lethality interaction with *iml3*Δ, which would normally encode an outer kinetochore protein required for pericentromeric cohesion during mitosis. This synthetic dosage lethality can be interpreted as a dominant negative effect of *XRS2* overexpression on DNA repair by nonhomologous end joining. The DNA repair interacts synergistically with DNA breaks likely associated with the loss of centromeric cohesion in the *iml3*Δ mutant. Indeed, *XRS2* overexpression is associated with slower growth, likely induced by genotoxic stress due to reduced nonhomologous end joining, which is suppressed by both *ddc1*Δ and *spt21*Δ mutations. *DDC1* encodes a DNA-damage checkpoint protein. Its absence is expected to prevent DNA damage checkpoint arrest in cells with reduced repair by nonhomologous end joining, and thus should restore growth rate at the expense of chromosome instability. This could explain the synthetic suppression interaction of *XRS2* overexpression with *spt21*Δ mutation, because *SPT21* abrogates transcription of two histone genes *HTA2-HTB2* and *HHF2-HHT2* that are required for chromatin-mediated signaling at broken DNA ends and triggering of DNA damage-induced checkpoint arrest.

Based these intuitive results, we expect that other matrices of genetic interactions derived form other experiments could reduce to a simple intuitive cycle. We have demonstrated that analyzing a matrix using integer linear programming to minimize the adjacent n-dimensional Euclidean distance results in a cyclical model that is both optimal and harmonious.

## Acknowledgements

We thank Eric Phizicky and Elizabeth Grayhack for supplying the MORF library; Phil Hieter for supplying strain *YPH275*; Raghavan Vasudevan, Vikashni Padmakumar, and Kankshit Bheda for transformations and plasmid purification; Arvind Kothandaraman, Rishov Chatterjee, and Hardeep Chiraya for laboratory logistics; Jose Salazar and Tanya Ferguson for Biomek FXP liquid handler support; Bryan Kraynack and Kirilynn Svay for training; Herbert Sauro, Alpan Raval, Craig Adams, Ali Nadim, and Susan Kane for discussions. This work was supported by grants from the National Science Foundation (#0527023, #0523643, #0523643 and #0941078), and the National Institutes of Health (# 1R01GM084881-01) to Animesh Ray. Part of the work was supported by the LANL National Flow Cytometry Resource (NFCR) funded by the National Center for Research Resources of NIH (Grant P41-RR01315). We thank Dr. Babetta L. Marrone (Director, NFCR) for helpful discussions on image analysis.

